# Molecular Atlas of the Adult Mouse Brain

**DOI:** 10.1101/784181

**Authors:** Cantin Ortiz, Jose Fernandez Navarro, Aleksandra Jurek, Antje Märtin, Joakim Lundeberg, Konstantinos Meletis

**Author notes:** these authors contributed equally (Cantin Ortiz, Jose Fernandez Navarro).

## Abstract

Brain maps are essential for integrating information and interpreting the structure-function relationship of circuits and behavior. We aimed to generate a systematic classification of the adult mouse brain organization based on unbiased extraction of spatially-defining features. Applying whole-brain spatial transcriptomics, we captured the gene expression signatures to define the spatial organization of molecularly discrete subregions. We found that the molecular code contained sufficiently detailed information to directly deduce the complex spatial organization of the brain. This unsupervised molecular classification revealed new area- and layer-specific subregions, for example in isocortex and hippocampus, and a new division of striatum. The whole-brain molecular atlas further supports the identification of the spatial origin of single neurons using their gene expression profile, and forms the foundation to define a minimal gene set - a brain palette – that is sufficient to spatially annotate the adult brain. In summary, we have established a new molecular atlas to formally define the identity of brain regions, and a molecular code for mapping and targeting of discrete neuroanatomical domains.

## Introduction

Mapping the adult brain, in terms of establishing reference maps of subregions and their borders, and determining the diversity of neuron types and their connectivity, is at the core of exploring the structure-function relationship of brain circuits that defines the diversity of animal behaviors *(1–4)*.

A central principle in generating brain atlases has over the past century been the annotation of tissue landmarks using microscopy to extract spatially relevant signals. These spatial definitions have to a large extent been based on cytoarchitectural features – such as differences in the density and form of cells – as well as chemoarchitectural definitions derived from distribution of key molecules such as neurotransmitters *(5, 6)*. Ongoing collective efforts in the field are now starting to reveal the details of neuron and region connections at the microscale and mesoscale level *(4, 7, 8)*, and gene expression in the human brain *(9, 10)*. These tissue definitions and the resulting mouse brain atlases *(11, 12)* have been essential for establishing the experimental framework to explore brain structure and function relationships, but have also resulted in debate and disagreement over the validity of expert-based region annotations *(13)*. Experimental neuroscience depends on the ability to repeatedly and accurately record and manipulate activity of neuron subtypes in specific brain regions, and genetic targeting approaches have therefore proven extremely valuable in cell-type specific targeting *(14, 15)*. Surprisingly, spatial classification has primarily been limited to layer-specific gene expression patterns in isocortex *(16)*, not capturing the great diversity of spatially segregated regions, such as the anteroposterior and mediolateral specialization in isocortex, or the more complex threedimensional organization of hippocampus and other subcortical regions. In fact, only a few brain regions have generally accepted border definitions, raising the importance of spatial definitions to describe nervous system organization from molecular patterns *(17)*.

We hypothesized that a molecular code could serve as a valuable mapping approach to define the spatial identity of brain subregions in an unbiased, systematic, and formalized fashion.

## Spatial transcriptomics at the whole-brain scale

We generated a whole-brain molecular atlas by capturing the spatial patterns of gene expression in the adult mouse brain using spatial transcriptomics (ST) *(18)*. We hybridized 75 coronal sections from one brain hemisphere covering the entire anteroposterior axis onto ST arrays (Fig. 1A, Fig. S1). Using a computational framework designed to generate reference maps *(19)*, we aligned each imaged brain hemisphere section, including the position of the ST spots in the tissue, with the Allen mouse brain reference atlas (ABA, http://www.brain-map.org/) *(12)* (Fig. 1B, Fig. S1). This established a complete brain atlas with information on tissue coordinates, the reference ABA neuroanatomical definitions, and spatially-defined gene expression patterns (Fig. 1C). We detected on average 4422 genes and 11.210 reads per spot (4422 +/-1164 unique genes, 11.210 +/− 5943 reads, Fig. S2). The complete data set after quality control contained information on the expression of 15.326 unique genes across 34.053 spots (Fig. 1D-E). As a first demonstration of the spatial expression patterns for well-established regional and cellular markers, we visualized the expression of the cortical marker Rasgrf2, the striatal marker Gpr88, and the thalamic marker Rora in 2D and 3D, and compared this to the ABA in situ hybridization (ISH) signal *(12)* (Fig. 1F). The correspondence between ISH patterns and the gene expression in the molecular atlas supported the possibility of using the complete molecular information in the atlas to extract patterns of spatial gene expression without any prior knowledge of spatial boundaries or regions.

**Fig. 1.**
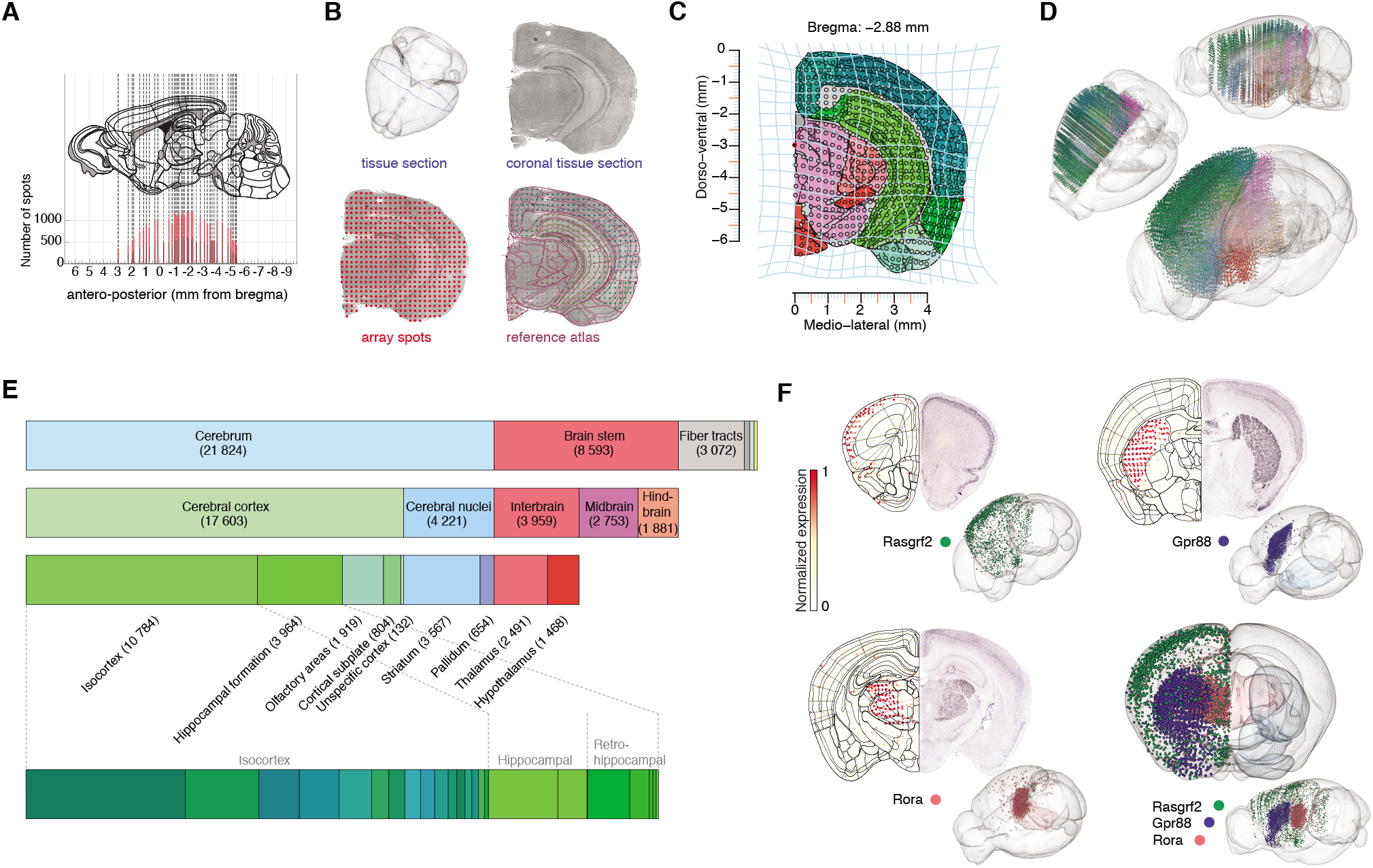
Spatial transcriptomics to generate a whole-brain molecular atlas. **(A)** Anteroposterior distribution of the coronal sections used to generate the molecular atlas (coordinates expressed in mm from bregma). Light red and dark red colors indicate adjacent sections. **(B)** Example of image processing pipeline (section at −2.88 mm posterior to bregma) showing the section outline in 3D (top left), the HE stained image (top right), the alignment of the HE image with the ST spots array (bottom left), and alignment with the mouse brain reference atlas (ABA, bottom right). **(C)** Transformation (light blue lines) of section shown in (B) to generate a common reference framework containing position of ST spots and neuroanatomical definitions. **(D)** Spatial mapping of all spots in 3D and color coded according to their regional identity. **(E)** Distribution of all spots according to main neuroanatomical region definitions. **(F)** Example of the tissue expression of three genes with spatially discrete signals in isocortex (Rasgrf2 in green), striatum (Gpr88 in blue), and thalamus (Rora in red). Left side: expression in the molecular atlas; right side: the ISH signal for each gene from the ABA. In 3D, right hemisphere shows spots with gene expression level above the 95th percentile. Left hemisphere shows region definition of striatum (light blue) and thalamus (light red). Composite 3D image showing expression of all three genes with regional definitions. Color scheme in (C), (D), (E), based on ABA reference color scheme.

## Computational approaches to extract new spatial domains from the molecular atlas

In order to generate a spatial classification of the adult mouse brain based exclusively on molecular patterns, we analyzed the ST gene expression data by performing dimensionality reduction and clustering. We performed an independent component analysis (ICAjade *(20)*) on top 50% variable genes to reduce the complexity of the dataset into a smaller number of dimensions. Each independent components (IC) represents a weighted combination of genes (gene loads). We visualized each IC in 2D and 3D in the reference atlas, allowing us to infer whether ICs contained spatial information (biological IC) or likely resulted from technical noise (technical IC) (Fig. 2A, and Fig. S3).

**Fig. 2.**
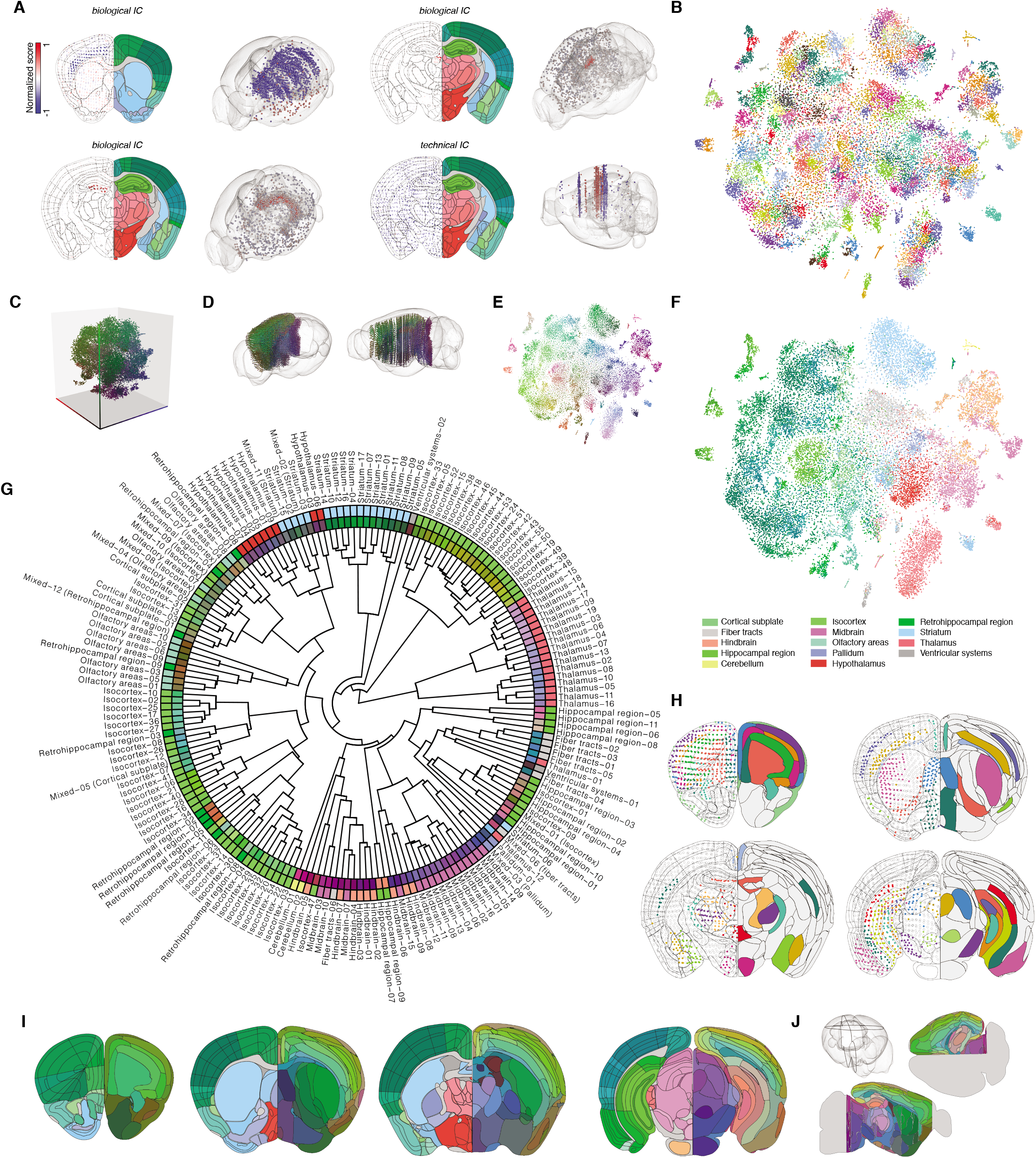
Computational approaches to extract new spatial domains from the molecular atlas. **(A)** Examples illustrating spatial expression of individual ICs (biological versus technical). Left side: IC signal (normalized score). Right side: corresponding ABA reference coronal section. IC scores above the 95th percentile in absolute value are shown in 3D. **(B)** t-SNE plot in 2D showing all individual spots and the 181 molecular clusters. Color scheme for each molecular cluster is based on a categorical color scheme (color palette of 27 unique colors that are repeated). **(C)** t-SNE plot in 3D showing all spots colored individually based on a RGB (red, green, blue) color space that corresponds to the three coordinates (x, y, z in the t-SNE space). **(D)** Spatial mapping of all spots in 3D colored by cluster similarity. Molecular clusters are color coded according to the RGB color space from (C) (RGB color based on median coordinates of spots per cluster for each of the three axes). **(E)** t-SNE plot in 2D showing all individual spots using same color scheme as in (D). **(F)** t-SNE plot in 2D showing all individual spots colored according to neuroanatomical subregions (ABA reference color scheme). **(G)** Hierarchical clustering of the 181 molecular clusters shown in a fan plot. Molecular clusters are named with a numerical identifier and a name according to spatial correspondence to subregion in ABA. The inner layer is color coded using molecular similarity color scheme as in (D) and (E). The outer layer is colored according to ABA subregions as in (F). **(H)** Examples of coronal sections from the molecular atlas. Left side: position and molecular identity of single spots belonging to selected clusters, and ABA subregion definitions in black lines. Right side: the same clusters shown in the molecular atlas (molecular atlas generated using SVM), black lines representing borders between molecular clusters. Color coded based on the categorical scheme used in (B). **(I)** Examples of coronal sections from the molecular atlas (right side, molecular clusters color coded based on molecular similarity as in (E)) compared to the ABA reference atlas (left side). **(J)** Visualization of virtual sectioning of the molecular atlas (shown in the glass brain using a black outline for two sections) in either a horizontal or a sagittal plane. Smoothed molecular clusters are shown with similarity color scheme in (E).

We selected 45 biological ICs to perform clustering *(21)*, resulting in 181 molecularly defined clusters. We visualized spots with a categorical color scheme according to their cluster identity in a whole-brain reference atlas (video S1) and in a 2D t-SNE *(22)* plot (Fig. 2B). We further generated a color scheme to visualize the molecular relationship between clusters by plotting spots in a 3D t-SNE plot, and coloring with a RGB (red, green, blue) color scheme corresponding to the x, y, z, coordinates (Fig. 2C). Plotting based on this color scheme showed the spatial distribution of molecularly similar clusters across the whole brain (Fig. 2D) as well as in a 2D t-SNE plot (Fig. 2E). Importantly, by applying a neuroanatomically relevant color code (the ABA reference color scheme) based on the known position of each spot in the atlas, clearly revealed the relationship between the molecular classification (based purely on the gene expression) and the broad spatial fingerprint of clusters (Fig. 2F, Fig. S4). To map the relationship between clusters, we used hierarchical clustering to generate a dendrogram for the 181 molecular clusters. We then color coded each cluster in a fan plot based on two definitions: the molecular similarity of clusters based on the 3D t-SNE RGB color code, and the color code denoting neuroanatomical identity (ABA reference color code, Fig. 2G). From these visualizations, we concluded that the molecular clustering respected spatial patterns at the global scale, and was aligned with certain aspects of current neuroanatomical definitions.

We next applied a support-vector machine (SVM) algorithm to transform the individual spots and corresponding molecular clusters into a continuous 3D atlas consisting of distinct molecular volumes (Fig. 2H, video S2). This resulted in a whole-brain atlas with detailed information on the spatial organization of molecular clusters (Fig. 2I, Fig. S5, video S3). The molecular atlas is openly available for use, further annotation or modification, and interactive visualization. The 3D molecular atlas further supports virtual sectioning of the brain at any angle (e.g. horizontal, sagittal) to easily visualize molecular clusters and their borders (Fig. 2J).

To explore the potential of molecular mapping as a road map for new spatial classifications at a finer scale, and to compare results from the molecular atlas with available reference maps, we first focused on the spatial organization of the isocortex. We used the neuroanatomical definition of isocortex and nomenclature of the ABA reference, which resulted in the annotation of 55 molecular clusters belonging to isocortex. Visualizing only these isocortical clusters in the t-SNE plot showed the underlying molecular-spatial structure that emerged purely from the gene expression patterns (Fig. 3A). The isocortical clusters showed clear separation by their layer identity, and to a large extent respected the established layer subdivisions, in terms of position as well as marker expression (Fig. 3B-D). More interestingly, we found that the molecular clustering also revealed the global organization of isocortical regions in the anteroposterior (AP) as well as mediolateral dimension (ML) (Fig. 3E). For example, molecular clusters were clearly segregated into distinct subregions of the frontal cortex (PL, ACA, ORB), forming borders between primary and secondary somatomotor and somatosensory cortex (Mop, MOs, SSp, SSs), and delineating spatially the retrosplenial cortex (RSP, Fig. 3F-I, videos S4). The molecular atlas did not differentiate between prelimbic and infralimbic parts of prefrontal cortex, but established borders between the medial prefrontal and orbital regions, including layer-specific definitions. In somatosensory isocortex, we found that the molecular clusters defined 8 separate layer-defining clusters, including molecular definitions of layer 5a, 5b, 6a, and 6b. Interestingly, we found that the molecular classification could separate layer 2/3 into two separate clusters with clear spatial organization. This layer 2/3 subdivision was evident in the somatosensory area (primary somatosensory, barrel field, SSp-bfd), as well as in other isocortical areas such as primary motor area (MOp). The hippocampal region (HIP) was subdivided into 11 molecular clusters, which spatially capture the unique hippocampal organization in 3D, and establish borders between CA1, CA2, and CA3. The dentate gyrus separated into three clear clusters that map onto the granule layer, molecular layer, and polymorph layer (i.e. hilus, Fig. S5). The molecular atlas of isocortex thereby supports detailed spatial classification of isocortex into discrete domains, that capture layer-specific subdivisions (dorsoventral axis, DV), together with spatial classification of the different isocortical subregion borders across the AP and ML axes.

**Fig. 3.**
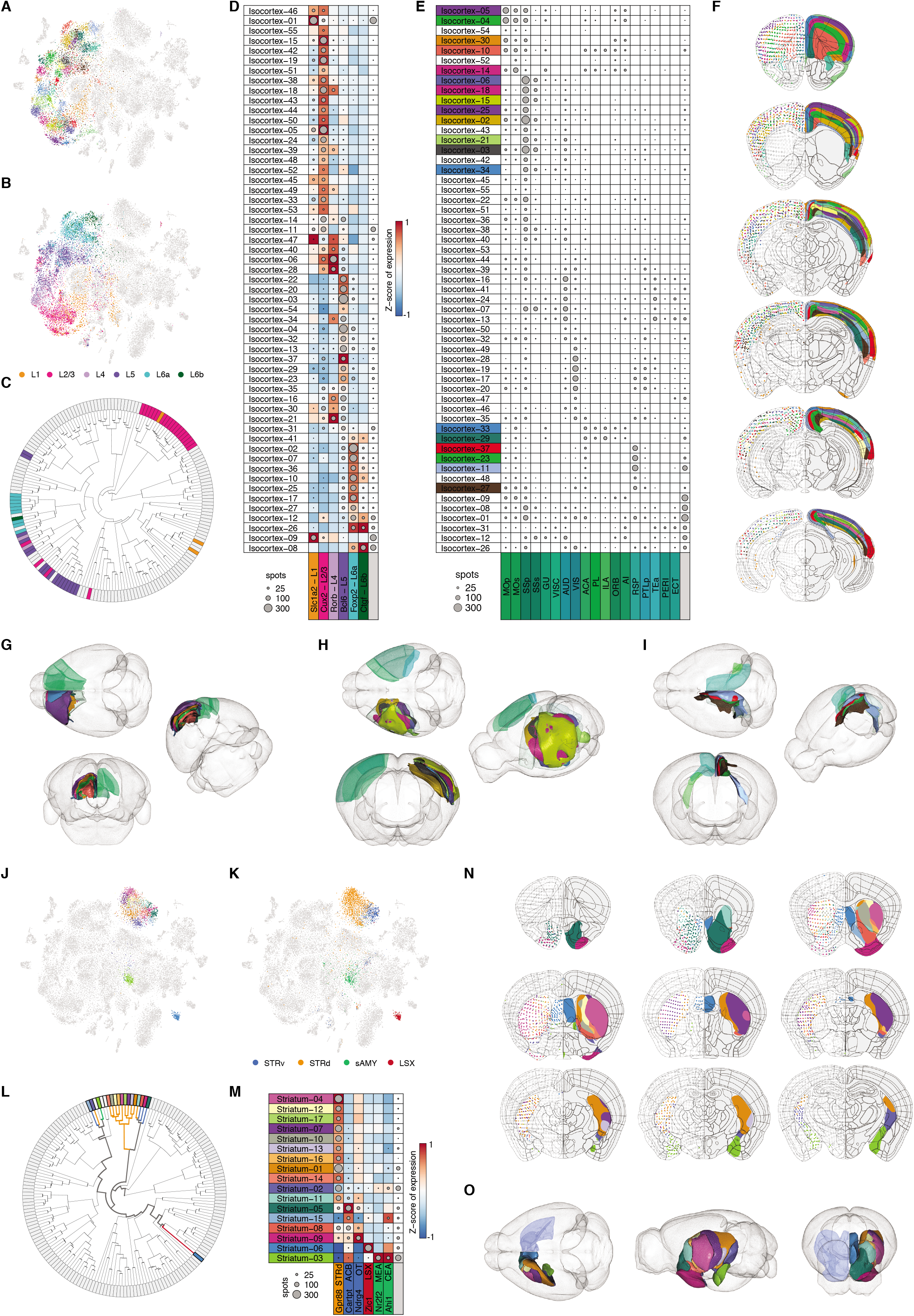
New molecular definitions of subregions and layers in isocortex and striatum. **(A)** t-SNE plot showing all spots belonging to isocortical clusters using the categorical color scheme. **(B)** t-SNE plot showing spots position across isocortical layers based on registration in the ABA. Color scheme represents layer identity. **(C)** Fan plot showing clusters according to their isocortical layer identity (color scheme as in (B)). **(D)** Visualization of all isocortical molecular clusters, and their gene expression pattern (averaged z-scored gene expression) for layer-enriched markers. Layer identity shown with same color code as in (B) and grey denoting non-cortical identity. **(E)** Visualization of all isocortical molecular clusters and their correspondence to major isocortical subregions (ABA color scheme, grey denoting non-cortical). Selected color-coded clusters are displayed in (G-I). **(F)** Visualization of all isocortical clusters in the molecular atlas in coronal sections across the anteroposterior axis. Left side shows individual spots belonging to molecular clusters. Right side shows the corresponding smoothed clusters in the molecular atlas. Black lines in both hemispheres show spatial definition of subregions based on the ABA. **(G-I)** Visualization in 3D of the molecular clusters found in different domains of isocortex. Left hemisphere shows clusters from the molecular atlas, and right hemisphere the corresponding subregions defined in the ABA. **(G)** Molecular clusters found in the anterior part of isocortex (PL, ILA, ACA, MO). **(H)** Molecular clusters found in the central part of isocortex (SS. MOp, PTLp). **(I)** Molecular clusters found in the medial and posterior part of isocortex (RSP, POST, PRE). **(J)** t-SNE plot showing all spots belonging to striatal clusters. **(K)** t-SNE plot showing all spots found in the striatum (STR) based on their spatial identity using the ABA definitions (STRv: ventral striatum; STRd: dorsal striatum; sAMY: striatumlike amygdala nuclei; LSX: lateral septal complex). **(L)** Molecular clusters corresponding to striatum shown in the hierarchical clustering. The dendrogram line coloring shows striatal subregions based on the ABA (same color scheme as (K)). **(M)** Visualization of all striatal molecular clusters, and their gene expression pattern (averaged z-scored gene expression) for selected spatial markers (same color scheme for regions as (K), and grey denoting non-striatal). **(N)** Visualization of all striatal clusters from the molecular atlas in coronal sections across the anteroposterior axis, using the same format as in (F). **(O)** Visualization in 3D of the molecular clusters covering striatal subregions. Left hemisphere shows molecular atlas and right hemisphere the striatum region (light blue, defined in the ABA). Dot surface in (D), (E), (M), corresponds to the absolute number of spots. Color scheme for molecular clusters in (A), (E-J), (L-O) based on the categorical color scheme.

To further explore how the molecular atlas can reveal spatial organization of subcortical areas, we focused on the striatum (STR), which in the ABA reference is divided into one major dorsal part of striatum (STRd), and the ventral striatum (STRv), including the lateral septal complex (LSX), and the striatum-like amygdalar nuclei (sAMY). Particularly the dorsal striatum has not been molecularly subdivided even if functional mapping suggests several discrete striatal subregions *(23)*. In the molecular atlas, STRd and the other striatal regions can be visualized in the t-SNE as 17 discrete clusters, and these molecular clusters reflect spatial information as shown when color-coded with the ABA reference color scheme (Fig. 3J-K). Importantly, these molecular clusters establish a new classification of the striatum into spatially-restricted subregions, including correlation with discrete marker expression (Fig. 3L-M). Visualization of the striatal clusters in 2D and 3D supports a spatial organization of dorsal striatum into several discrete molecular domains across the AP and ML axes, in addition to the ventral and septal divisions (Fig. 3N-O, video S5).

In summary, capturing the spatial gene expression signatures of the adult mouse brain allowed us to produce a detailed whole-brain atlas built from only molecular information, thereby establishing a formal system for spatial definitions in neuroanatomy.

## Mapping the spatial origin of single cells using the molecular atlas

The molecular profiling of neurons based on single-cell RNA sequencing (scRNA-seq) has been used to establish a molecular code for classification of neuron subtypes. This classification scheme usually lacks information on the spatial origin of the classified neurons. We investigated whether the molecular atlas could serve as a resource to map the spatial origin of single neurons based on their scRNA-seq molecular profiles, thereby providing a spatial dimension to the cell-type definitions in large-scale cell classification efforts. To provide proof of principle of the spatial mapping approach, we used available data *(24)* on single-cell RNA-seq of 23.822 neurons and non-neuronal cells isolated from either primary visual cortex (VISp) or from the frontal cortex (anterior lateral motor cortex, ALM).

We trained a neural network classifier on the molecular atlas to develop predictions on the spatial origin of each single cell, and quantified spatial mapping onto molecular clusters in ALM and VISp (Fig. 4A-B). When we used a cut-off in the prediction probability to select 50% of the neurons with the highest confidence in spatial mapping (0.99 probability), we found that 76% of the glutamatergic neurons were accurately classified onto the correct region (VISp versus ALM, 65% without thresholding) and 37% to the correct region and layer (Fig. 4C-D, Fig. S6). In contrast, we found that GABAergic neurons, as well as the different non-neuronal cells, contained limited information about spatial origin and could not be accurately mapped onto the molecular atlas (Fig. 4E). In summary, this demonstrates the value of the molecular atlas as a resource to map the spatial origin of neurons using single-cell molecular information.

**Fig. 4.**
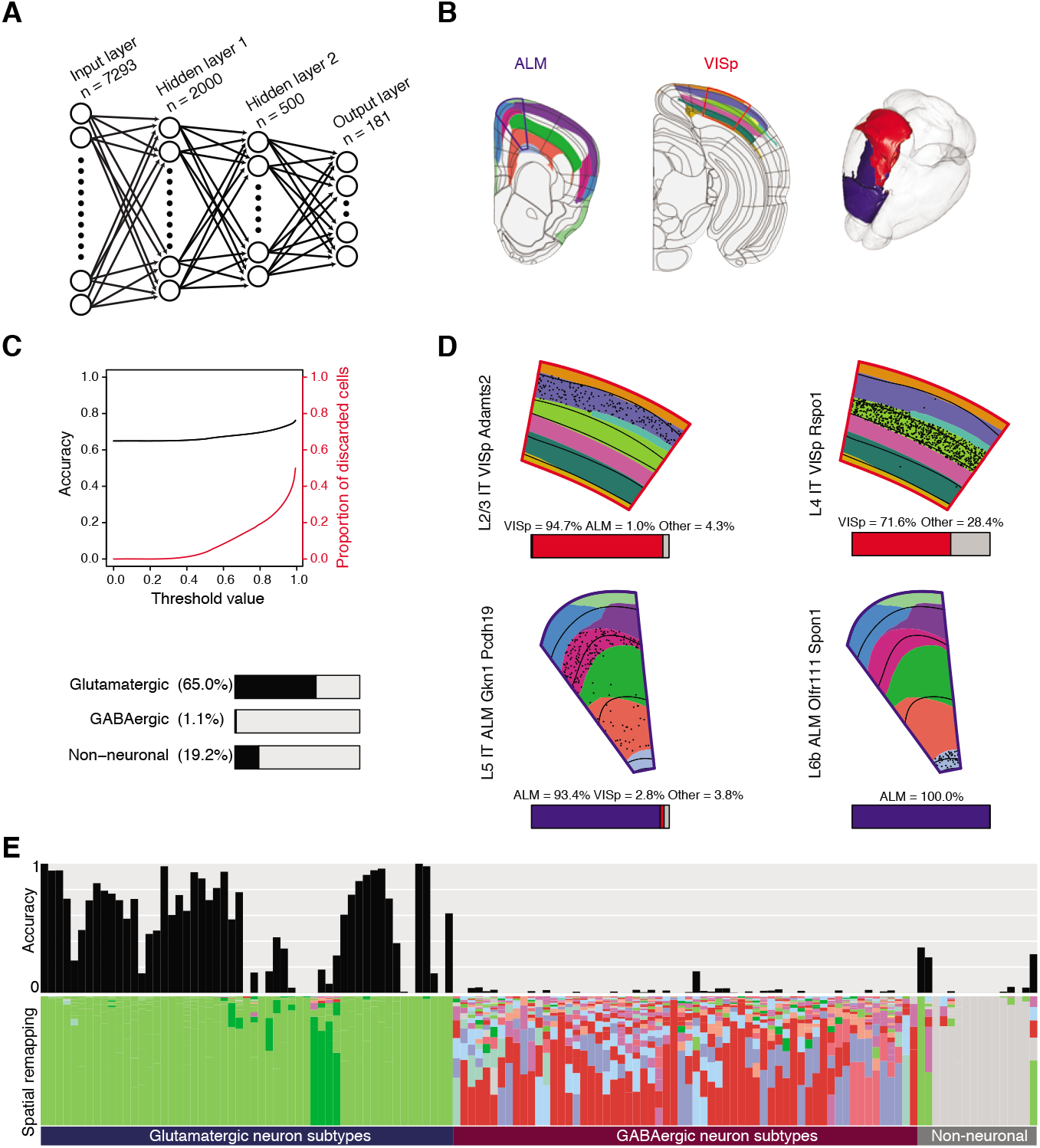
Mapping the spatial origin of single cells using the molecular atlas. **(A)** Schematic of the neural network (NN) organization. **(B)** Showing clusters from the molecular atlas that correspond to ALM (blue line) and VISp (red line). Black lines show spatial definitions from the ABA reference atlas. Right side: 3D visualization of the clusters corresponding to ALM (blue) and VISp (red). **(C)** Top: spatial mapping accuracy for single neurons (glutamatergic) and discarded cells as a function of the probability threshold value. Bottom: accuracy of spatial mapping of single cells for the three main cell types. **(D)** Visualization of the spatial mapping of single cells from distinct scRNA-seq clusters isolated from VISp (red outline) or ALM (blue outline). Spatial mapping of individual neurons in scRNA-seq clusters belonging to VISp (top row) and ALM (bottom row). Molecular atlas clustering shown with the categorical color scheme and layer definitions shown with back lines (ABA reference). Black dots represent spatial mapping of single neurons onto predicted cluster. Quantification of spatial mapping region is also shown for each scRNA-seq cluster with stacked bar plot. **(E)** Top row shows the accuracy of the spatial mapping of all single cells using the NN predictions for each discrete scRNA-seq clusters (133 clusters, 23.822 cells). The order of cell types is based on the published hierarchical clustering, showing major cellular identity (glutamatergic, GABAergic, non-neuronal). Bottom row shows the proportion of mapped cells for each scRNA-seq cluster onto the molecular clusters. Color scheme of molecular clusters based on the ABA reference colors.

## A reduced brain palette captures whole-brain organization

We aimed to determine a reduced gene set that would be sufficient to capture at the global scale the major brain subdivisions. The complete molecular atlas is based on the gene expression patterns of 7663 genes. In order to establish in an unsupervised fashion the genes that best capture global spatial signals, we selected genes from either the ICs or from weights in a SVM model, and used the normalized mutual information index (NMI) as a measurement of cluster similarity (Fig. 5A). A panel of 266 unique genes was established on the basis of a NMI 0.6 for clustering using genes from the ICs. We found that the 266 genes – forming a brain palette – when used to define the brain into 181 molecular clusters, established a whole-brain spatial division similar to the clustering in the full molecular atlas (Fig. 5B-D). We calculated the cluster similarity between the molecular atlas and the reduced version in order to visualized the pairwise correlation between spatially conserved subregions (Fig. 5E, Fig. S7). We found that the spatial definitions of molecular domains were conserved globally (Fig. 5F), and as shown for example in a cluster group consisting of several isocortical, retrohippocampal, and olfactory area subregions (Fig. 5G-H), and in the main hippocampal clusters (Fig. 5I). We concluded from this analysis, that molecular clusters and large domains are well-defined spatially based on the reduced brain palette, whereas some regions probably require a richer gene expression dataset for detailed subdivision (Fig. 5, Fig. S8).

**Fig. 5.**
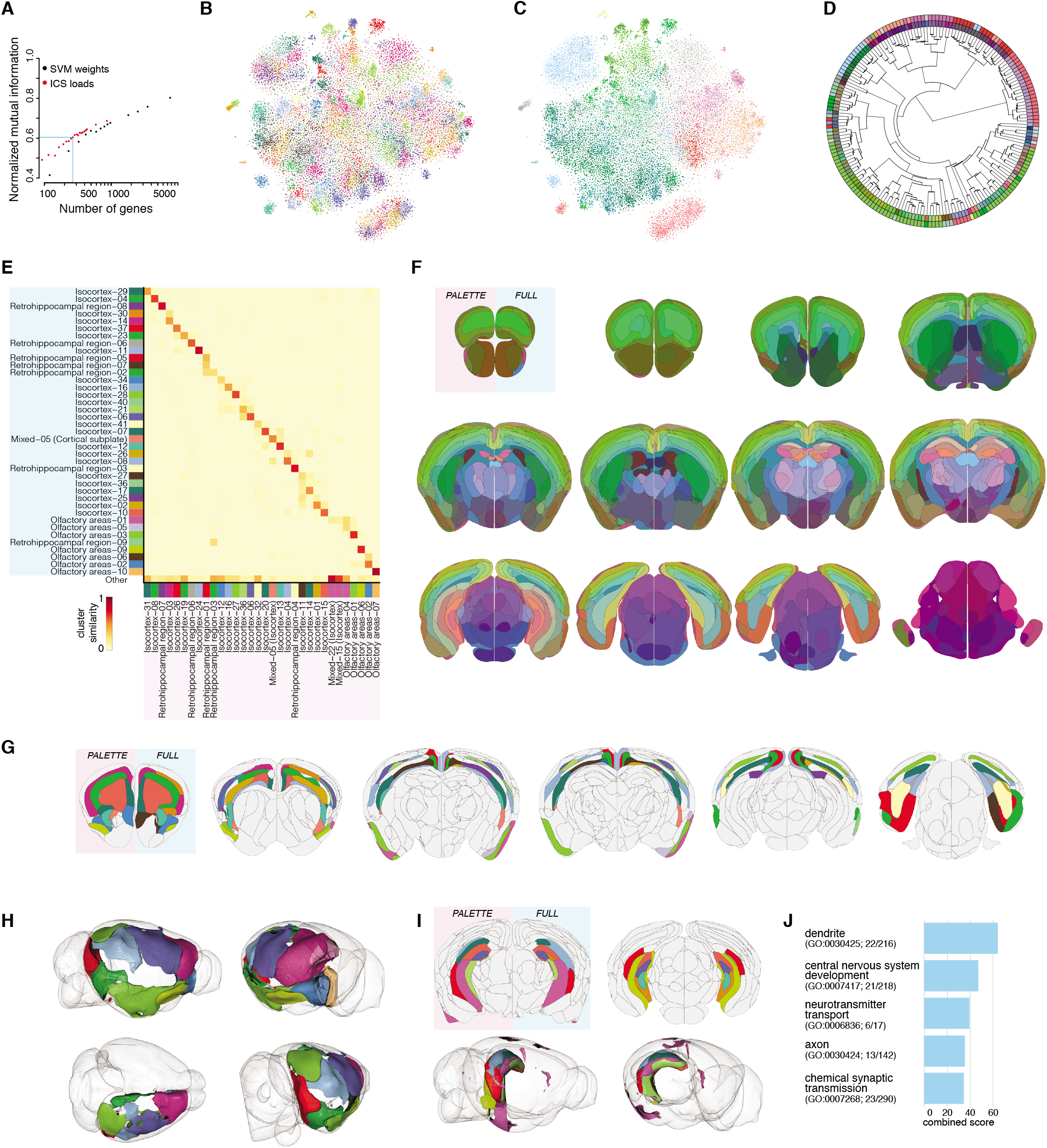
A reduced brain palette captures the whole-brain spatial organization. **(A)** The normalized mutual information (NMI) used to select the gene set to establish the brain palette. **(B)** t-SNE plot showing 181 molecular clusters based on the brain palette (266 genes). Clusters are colored based on the best matching cluster in the complete molecular atlas using the same categorical color scheme. **(C)** t-SNE plot in 2D showing all individual spots colored according to neuroanatomical subregions (ABA reference color scheme). **(D)** Hierarchical clustering of the brain palette 181 molecular clusters shown in a fan plot. The inner layer is color coded using molecular similarity color scheme. The outer layer is colored according to ABA subregions as in (C). **(E)** Example of the similarity between the full molecular atlas (y axis) and the brain palette molecular atlas (x axis) for selected clusters. Categorical color scheme used to visualize the same clusters in (G) and (H). Fig. S7 shows the entire similarity matrix. **(F)** Visualization of the brain palette molecular atlas (left side, pink background in the first section) and the full molecular atlas (right side, blue background in the first section) based on molecular similarity color scheme. **(G)** Visualization of isocortical, retrohippocampal and olfactory clusters displayed in (E) with categorical color scheme from (B) from the brain palette molecular atlas (left side, pink background in the first section) and the full molecular atlas (right side, blue background in the first section). **(H)** Visualization in 3D of selected isocortical clusters from the brain palette molecular atlas with categorical color scheme from (B). **(I)** Comparison of hippocampal clusters from the brain palette molecular atlas (2D: left side with pink background in the first section, 3D: right hemisphere) and the full molecular atlas (only shown in 2D: right side with light blue background in the first section), categorical color scheme from (B). **(J)** Top five significant enriched GO terms for genes in the brain palette (266 genes).

In order to establish whether the composition of the brain palette could reveal something about the underlying biological mechanisms that establish or maintain spatial definitions, we analyzed the brain palette genes based on their assignment to gene ontology (GO) terms. Overall, we found significant enrichment for 64/266 genes, belonging to five GO families (combined score > 30, adjusted p value <1e-3, Fig. 5J, Fig. S9). These genes are associated with cellular processes (dendrite, axon), and biological processes (nervous system development, neurotransmitter transport and synaptic transmission). The enrichment in genes involved in nervous system development, as well as molecular aspects of dendrite and axon compartments, points to the key biological processes that establish brain regionalization, form and possibly maintain spatial patterns in the adult brain. The minimal version of the molecular atlas builds on the brain palette – a list of genes whose expression is sufficient to recapitulate most aspects of the spatial whole-brain subdivisions – revealing some of the biological processes behind spatial identity, while also forming an important experimental tool for mapping global brain space with a small number of genes.

## Discussion

We have here established a molecular atlas of the adult mouse brain exclusively on the basis of unsupervised classification of spatial transcriptomic patterns on a whole-brain scale. A challenge in neuroscience has for long been the formalized definitions of functionally and molecularly relevant subdivisions of the nervous system. The molecular definition of brain regions is an unbiased approach to map the discrete spatial domains at the whole-brain scale, and importantly also introduces the ability to develop molecular approaches with spatial information to study nervous system organization, the evolutionary conservation of molecular and spatial codes, and their relationship to physiology and pathophysiology.

We found that it is possible to develop a detailed whole-brain spatial annotation based purely on gene expression signatures, for well-studied regions of the brain such as the somatosensory and somatomotor area in isocortex, and the hippocampus, as well as for major regions with poorly defined subdivisions such as the dorsal striatum. In addition, the molecular atlas also establishes the identity of isocortical layers across the entire isocortex, olfactory areas (e.g. piriform area), and the retrohippocampal area (e.g. entorhinal area). The subdivision of these areas into specific layers, and how layer divisions are conserved across different areas has remained unclear and debated *(25)*. We here propose a molecular definition of layers that contains subregional information, and for example provide evidence for discrete molecular separation of layer 2/3. It remains to be determined whether our inability to identify clear subdivision in for example parts of the prefrontal cortex *(26)*, or in small amygdalar areas, is due to the absence of molecular identifiers or is a result of another division level (e.g. connectivity *(27)*).

Our work furthermore defines a reduced gene set – a brain palette – that is sufficient to capture the spatial complexity of brain subregions on a whole-brain scale. We have demonstrated the power of region classification based on a reduced brain palette, and this approach can be similarly used to define any region of interest, by developing local palettes that define subregions and borders with increased spatial resolution. Similar to the classification of neuron subtypes, which is often defined by combinatorial gene expression rather than single markers, the molecular classification of regions and borders will to some extent depend on combinatorial expression of some genes. In that sense, the brain palette establishes a reduced set of genes that are sufficient to map the entire adult brain into the relevant subregions. Thereby, the molecular atlas is the basis for also establishing a specific regional palette for any region of interest, to guide spatial annotation of subregions and borders using existing methods to map gene expression in tissue *(28–33)*. We anticipate that technological advancements will lead to increase spatial and sequencing resolution *(34, 35)*, potentially also leading to a more detailed molecular classification of cell types and brain regions.

We do not know the function of the molecules that define spatial subdivisions, but we provide evidence that certain key gene ontologies (related to dendrite and axon processes, nervous system development, and glutamatergic neurotransmission) are probably major contributors to the spatial classification in the adult brain. The signals that underlie global division of the brain into subregions is associated with developmental relevant genes, illustrating the role of the developmental axes as important determinants of the adult regionalization. Previous work has shown a relationship between gene expression in the adult brain and embryonic development in rodents *(36)* and humans *(37)*, supporting the role of developmental trajectories in brain maps. However, the mechanisms whereby the gene products contribute to the function or maintenance of regional identity in the adult remain unknown. A key question for the future will be to what extent the molecular identifiers of regionalization and spatial domains are evolutionary conserved, similar to conservation of cell markers across species *(38)*. For example, how is cortical expansion in the primate brain a result of an expansion of conserved molecular domains found in other vertebrates, or does possible new function arise from new spatiomolecular definitions. It will therefore be important to map the molecular signatures of regionalization in other species, including humans, and to thereby determine the conserved brain palette.

Classification of the adult brain from a molecular perspective should eventually be fused with large-scale connectivity and activity maps, to produce structure-function maps of regions and circuits. Ultimately, generating whole-brain molecular atlases in mouse models of brain disease, and comparing the conserved molecular signatures that define brain regionalization across species, can serve as key points for developing a comprehensive understanding of the role of circuits in normal behavior, and provide templates for new strategies to treat brain disorders.

## Acknowledgments

The authors thank Jonas Frisén, Daniel Fürth, Annelie Mollbrink, Emil Wärnberg, Pierre Le Merre, Hans Brünner, Konstantinos Ampatzis, Marie Carlén, for providing advice and technical help.

This study was supported by the Swedish Research Council (VR 2017-01457 to K.M.), the Swedish Brain Foundation (Hjärnfonden grant to K.M.), the Swedish Foundation for Strategic Research (SSF grant FFL12-0006 to K.M.), the Knut and Alice Wallenberg Foundation (KAW grant to J.L.), the Olav Thon Foundation (doctoral and postdoctoral funding for J.F.N. and A.J., respectively), EU JPND INSTALZ (doctoral and postdoctoral funding for J.F.N. and A.J., respectively), Karolinska Institutet (senior research fellow grant to K.M., doctoral funding KID grant to A.M.).

The supplementary materials contain additional data.

